# Early selection of the amino acid alphabet was adaptively shaped by biophysical constraints of foldability

**DOI:** 10.1101/2022.06.14.495995

**Authors:** Mikhail Makarov, Alma C. Sanchez Rocha, Robin Krystufek, Ivan Cherepashuk, Volha Dzmitruk, Tatsiana Charnavets, Anneliese M. Faustino, Michal Lebl, Kosuke Fujishima, Stephen D. Fried, Klara Hlouchova

## Abstract

Whereas modern proteins rely on a quasi-universal repertoire of 20 canonical amino acids (AAs), numerous lines of evidence suggest that ancient proteins relied on a limited alphabet of 10 ‘early’ AAs, and that the 10 ‘late’ AAs were products of biosynthetic pathways. However, many non-proteinogenic AAs were also prebiotically available, which begs two fundamental questions: Why do we have the current modern amino acid alphabet, and Would proteins be able to fold into globular structures as well if different amino acids comprised the genetic code? Here, we experimentally evaluated the solubility and secondary structure propensities of several prebiotically relevant amino acids in the context of synthetic combinatorial 25-mer peptide libraries. The most prebiotically abundant linear aliphatic and basic residues were incorporated along with or in place of other early amino acids to explore these alternative sequence spaces. We show that foldability was a critical factor in the selection of the canonical alphabet. Unbranched aliphatic and short-chain basic amino acids were purged from the proteinogenic alphabet despite their high prebiotic abundance because they generate polypeptides that are over-solubilized and have low packing efficiency. Surprisingly, we find that the inclusion of a short-chain basic amino acid also decreases polypeptides’ secondary structure potential. Our results support the view that despite lacking basic residues, the early canonical alphabet was remarkably adaptive at supporting protein folding and explain why basic residues were only incorporated at a later stage of the alphabet evolution.

## INTRODUCTION

The role of peptides and proteins in life’s emergence has historically been sidelined as they have been, at times, perceived as only relevant after the evolution of nucleotide-based polymers, the genetic code, and a translation apparatus. However, the prebiotic abundance of amino acids and their ease of condensation resulting in polymers capable of creating functional hubs along with various cofactors (such as metal ions and organic compounds) has prompted some chemists to reconsider peptides’ role during life’s early evolution.^1–4^ While extant proteins are built from a sequence space spanned by the twenty canonical amino acids (cAAs) and rely on the specificity of the Central Dogma, early peptides (and peptide-like polymers) likely arose from a larger pool of prebiotically plausible monomers.

The modern protein alphabet was most likely selected during the first 10–15% of Earth history (~4.5–3.9 Ga) but the factors guiding its chemical and biological evolution remain unclear.^5^ Some of the cAAs are thought to be “late” additions to the genetic code, enabled only (or enriched significantly) by the evolution of their biosynthetic pathways. Indeed, many of the late amino acids are considered to be scarce in the prebiotic synthesis experiments.^6^ “The “early” amino acids (Ala, Asp, Glu, Gly, Ile, Leu, Pro, Ser, Thr and Val) were apparently available through prebiotic synthesis along with many other non-canonical amino acids (ncAAs) and their alternatives (such as beta- and gamma-amino acids, hydroxy acids or dicarboxylic acids). Based on two independent meta-analyses and many parallel studies, the canonical alphabet was therefore selected first from the large pool of prebiotically plausible structures and then supplemented with late amino acids through metabolism.^7,8^ Why certain amino acids were selected over others (and when) has been repeatedly discussed in literature over the last few decades.^5,9–12^ This question can be further subdivided into two separate but related questions, namely: (i) Why were the ten early cAAs selected from the prebiotic environment, and (ii) What factors drove the selection of the additional residues in the following era? Has protein evolution been successful as a consequence of the selected cAAs, or could similar protein space be formed with alternative alphabets?^9^

While some of the choices for the early amino acids are quite expected (especially for the C2 and C3 amino acids), others have been repeatedly considered thought-provoking. Most strikingly, the early amino acid alphabet lacked positively charged residues, even though diaminopropionic and diaminobutyric acids (C3 and C4 amino acids, DAP and DAB, respectively) have been shown to be prebiotically plausible.^13,14^ Neither lysine nor arginine are considered prebiotically plausible, whilst ornithine is known to cyclize, promoting peptide chain scission.^9^ Moreover, the canonical alphabet does not include some of the most abundant linear aliphatic amino acids such as α-amino-n-butyric acid (ABA), norvaline (Nva) and norleucine (Nle) (C4, C5, and C6, respectively) while their branched alternatives (such as Val, Leu, Ile) were selected^9^ despite being available at much reduced abundances. It has been hypothesized that perhaps the earlier version of the amino acid alphabet included some ncAAs that were later replaced by late cAAs.^2,15^ This is an intriguing hypothesis especially in light of the fact that some of these ncAAs have been reported as promiscuous substrates of some aminoacyl-tRNA synthetases.^16^

This study experimentally determines the properties that would have accompanied some of the most feasible ncAA candidates from the prebiotic pool. We incorporated the selected ncAAs into combinatorial peptide libraries along with (or replacing) other early amino acids to evaluate their effect on fundamental physicochemical properties such as solubility and secondary structure formation. The outcomes of our study show that an inclusion of some of the most feasible alternatives would *not* have supported the twin requirements of rich secondary structure potential and solubility. Hence, our study demonstrates that biophysical constraints on foldability provided a selective pressure shaping the building block selection in the early alphabet and therefore explains why certain early AAs (and not others) were selected to construct proteins.

## RESULTS

### Design and synthesis of peptide libraries from alternative alphabets

25-mer combinatorial peptide libraries were synthesized using solid-support chemical synthesis with isokinetic mixtures of amino acids (see Supplementary Table S1).

To model the sequence space available to different subsets of the amino acid alphabets, the libraries included the entire canonical alphabet without Cys (19F; F = full), its prebiotically available subset of 10 (10E; E = early), an alternative of 10E where the branched aliphatic amino acids were substituted with their unbranched prebiotically-abundant alternatives (10U; U = unbranched), the 10E library supplemented with: Arg as a representative of a modern cationic cAA (11R; R = Arg); or DAB as a representative of a potentially early cationic AA (11D; D = 2,4-diaminobutyric acid); or Tyr as a representative of an aromatic AA (11Y; Y = Tyr) (Figure 1).

**Figure 1.**
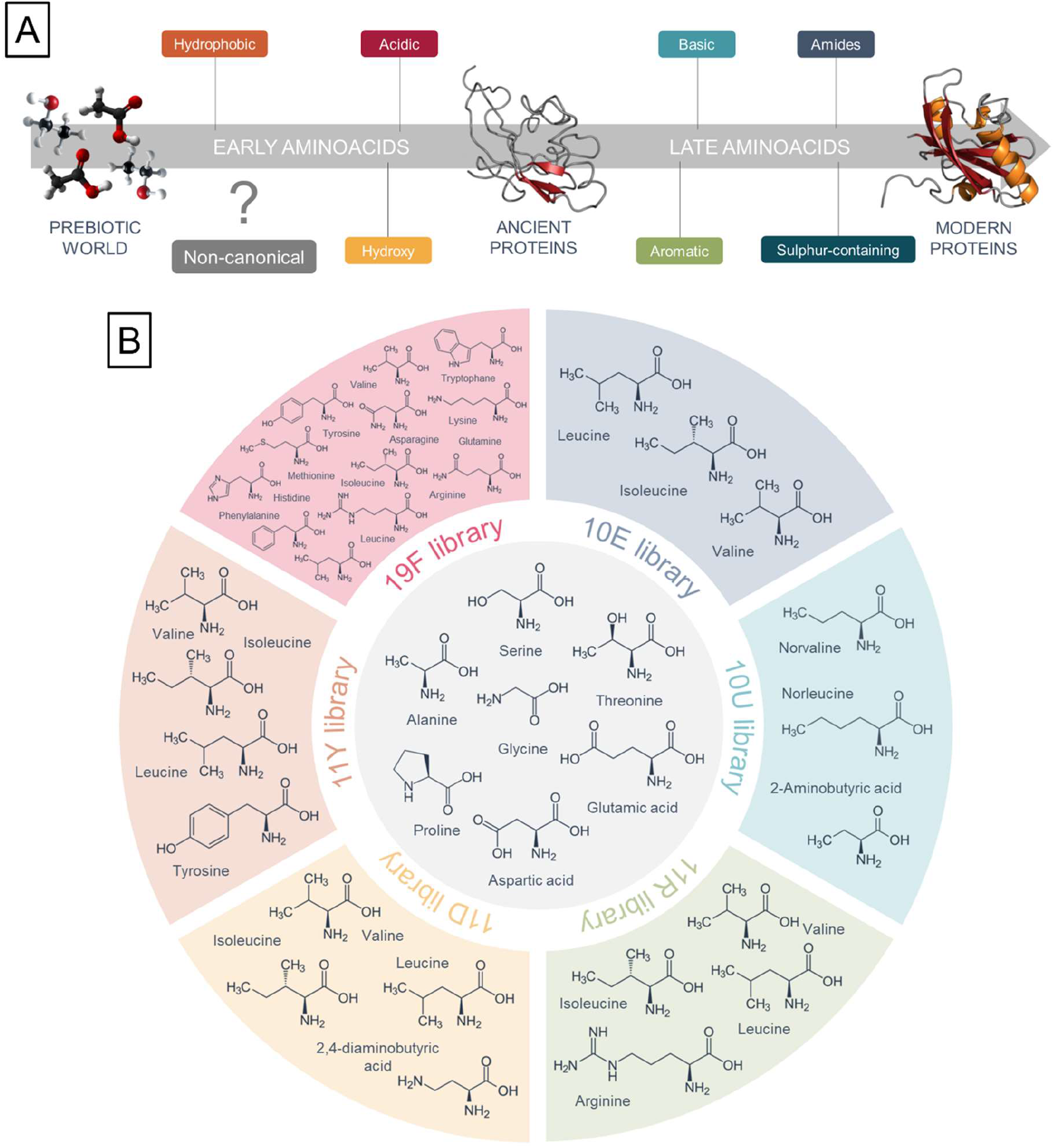
The assumed early and late stages of amino acid alphabet incorporation during protein evolution (A) and design of peptide libraries based on this order (B).

MALDI spectra confirmed the expected molecular weight range and distribution of the combinatorial libraries, reflecting their respective compositions (Supplementary Figure S1). Apart from the DAB in 11D library, the amino acid analyses of all libraries confirmed the expected composition (Supplementary Figure S2). The analysis protocol could not detect DAB whose presence in the 11D library (contrary to the 10E library) was instead determined using fluorescamine assay (Supplementary Figure S3). All the libraries passed quality control and were lyophilized to establish Cl^-^ as the counterion for all downstream analyses.

### Solubility and aggregation profiles

The ability to remain soluble under different conditions could represent an important selection factor during the formation of the early protein alphabet, as prebiotically relevant environments spanned from alkaline hydrothermal vents to acidic lakes.^17,18^ We measured the solubility profiles of these peptide libraries in the pH range 3–11, and also at low vs. high ionic strength, spectrophotometrically by absorption of peptide bonds at 215 nm and fluorometrically in case of library 19F due to its very poor solubility (Figure 2). In general, ionic strength did not significantly modulate solubility, with the exception of the 11Y library (see below).

**Figure 2.**
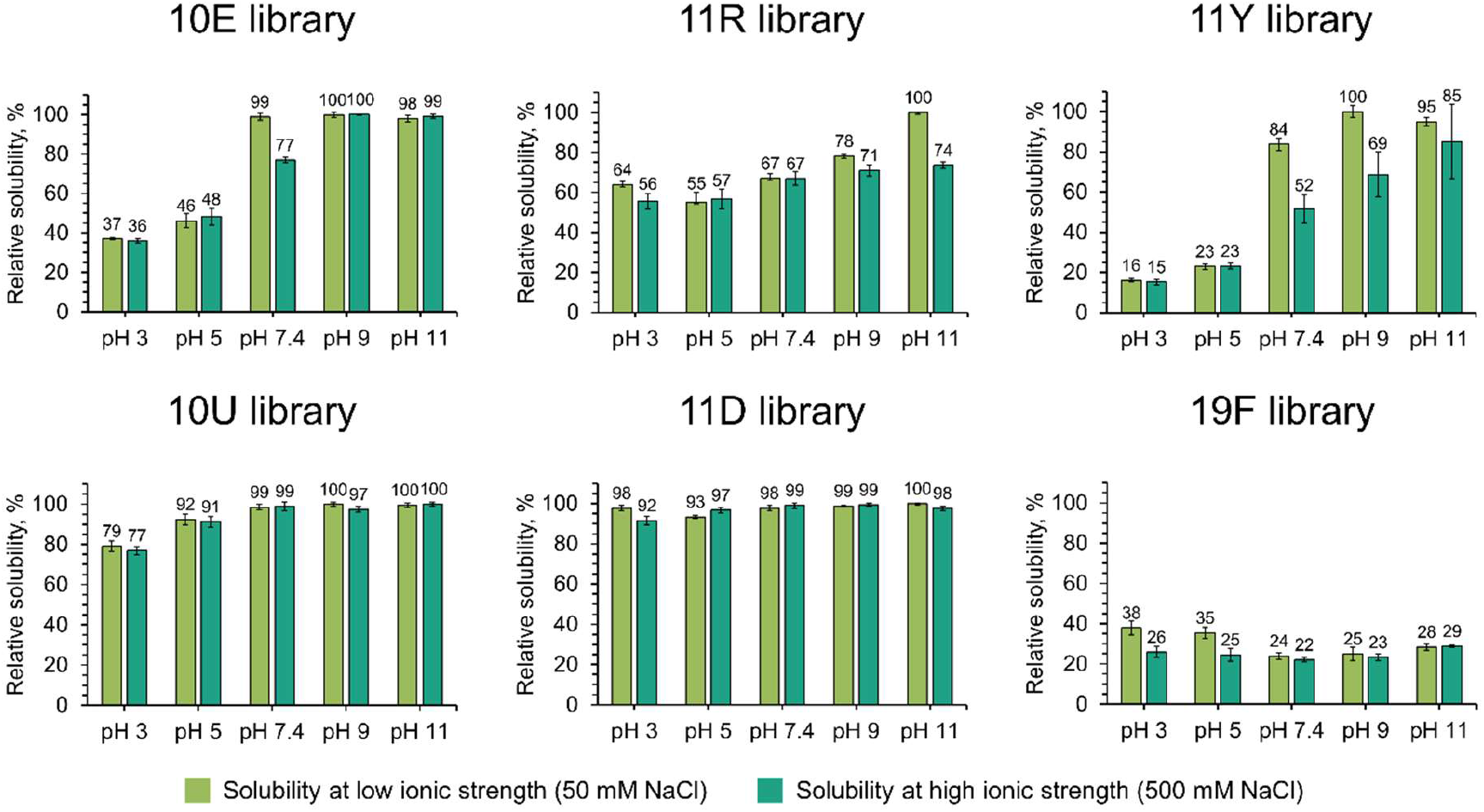
Solubility of 25-mer combinatorial peptide libraries at different pH and ionic strengths. Solubility was measured for 0.5 mg/ml nominal peptide library solutions in 20 mM ABP buffers (pH 3–11) at 50 and 500 mM NaCl. Solubility of 19F peptide library was measured by fluorescence spectroscopy, and solubility of the other peptide libraries was measured by UV-Vis absorption spectroscopy.

At a fixed nominal peptide concentration (0.5 mg/ml), the 19F library is significantly less soluble than the libraries with narrower amino acid repertoires. This result is in line with previous studies reporting that subsets of the canonical library that are enriched with the early amino acids are significantly more soluble than the full alphabet version.^19–21^ This phenomenon could be partly explained by the physicochemical nature of the full alphabet (such as the presence of aromatic amino acids) leading to increased aggregation propensity,^22^ or the greater ‘search complexity’ of a larger alphabet (leading to a more rugged landscape). Regardless of the mechanism, the results support the view that peptides composed of the full alphabet were more reliant on chaperones and/or translation to fold.^2,22^

The solubilities of the canonical library subsets (10E, 11R and 11Y) trend upwards with increasing pH (going from pH 3 to pH 11) and 11Y (which includes Tyr as a representative aromatic amino acid) is the least soluble of these three as expected. This result can be potentially explained by pointing out that all of these alphabets (as do modern proteomes)^23^ tilt toward being acidic; hence, at alkaline pH the peptides will accrue larger net negative charge and will repel one another electrostatically (inhibiting aggregation). This is broadly consistent with the practice of using mildly-alkaline buffers in protein refolding experiments, which tend to facilitate refolding by preventing aggregation.^24^

The two libraries that include the non-canonical alternatives of aliphatic (10U) and basic (11D) amino acids are strikingly more soluble than 10E and 11R throughout the examined pH range (Figure 2). Both of these results are unexpected for distinct reasons. Based on the physical chemistry of elementary hydrocarbons, one might expect that the 10U library would be less soluble because linear hydrocarbons are more hydrophobic than branched hydrocarbons of equal carbon-count because they create cavities with greater surface area.^25^ Moreover, the inclusion of a basic amino acid would raise the average isoelectric point of the library (relative to 10E) to be closer to neutral, which would decrease intermolecular repulsion, and thereby increase aggregation.

The soluble fractions of the libraries were additionally screened for the occurrence of soluble aggregates using size-exclusion chromatography (Supplementary Figures S4 and S5, in low and high ionic strength, respectively). No significant amount of such phenomenon was observed with the exception of 11Y and 10E libraries where a minor (up to ~10%) fraction of soluble aggregates was detected (Supplementary Figure S4). Overall, the elution profiles correlated with the spectrophotometric solubility measurements in which the 11R library was the most soluble of the canonical subset libraries, and the 10U and 11D libraries were fully soluble across the entire pH range.

### Secondary structure propensity

Natural non-folding sequences as well as random sequences with low secondary structure content have been previously reported to be highly soluble.^21,26^ The solubility profiles of the 11D and 10U libraries therefore suggest that the incorporation of the respective ncAAs may decrease the potential for secondary structure formation. CD spectroscopy was used to estimate average secondary structure content of the libraries over the studied pH range (Figure 3) and upon its induction with 2,2,2-trifluoroethanol (TFE) in pH 7.4 (Figure 4). Because no dominant secondary structure motif is expected in these highly diverse libraries and we are rather interested in average structural propensities, the secondary structure content was evaluated as the ratio of CD signal at 222 nm to the CD signal at 200 nm (shown by inlet graphs in Figures 3 and 4), because both alpha helices and beta sheets exhibit higher ellipticities than random coils at 222 nm.^27^ We did not observe any significant induction of secondary structure upon changing the metal ion concentration (Supplementary Figure 6).

**Figure 3.**
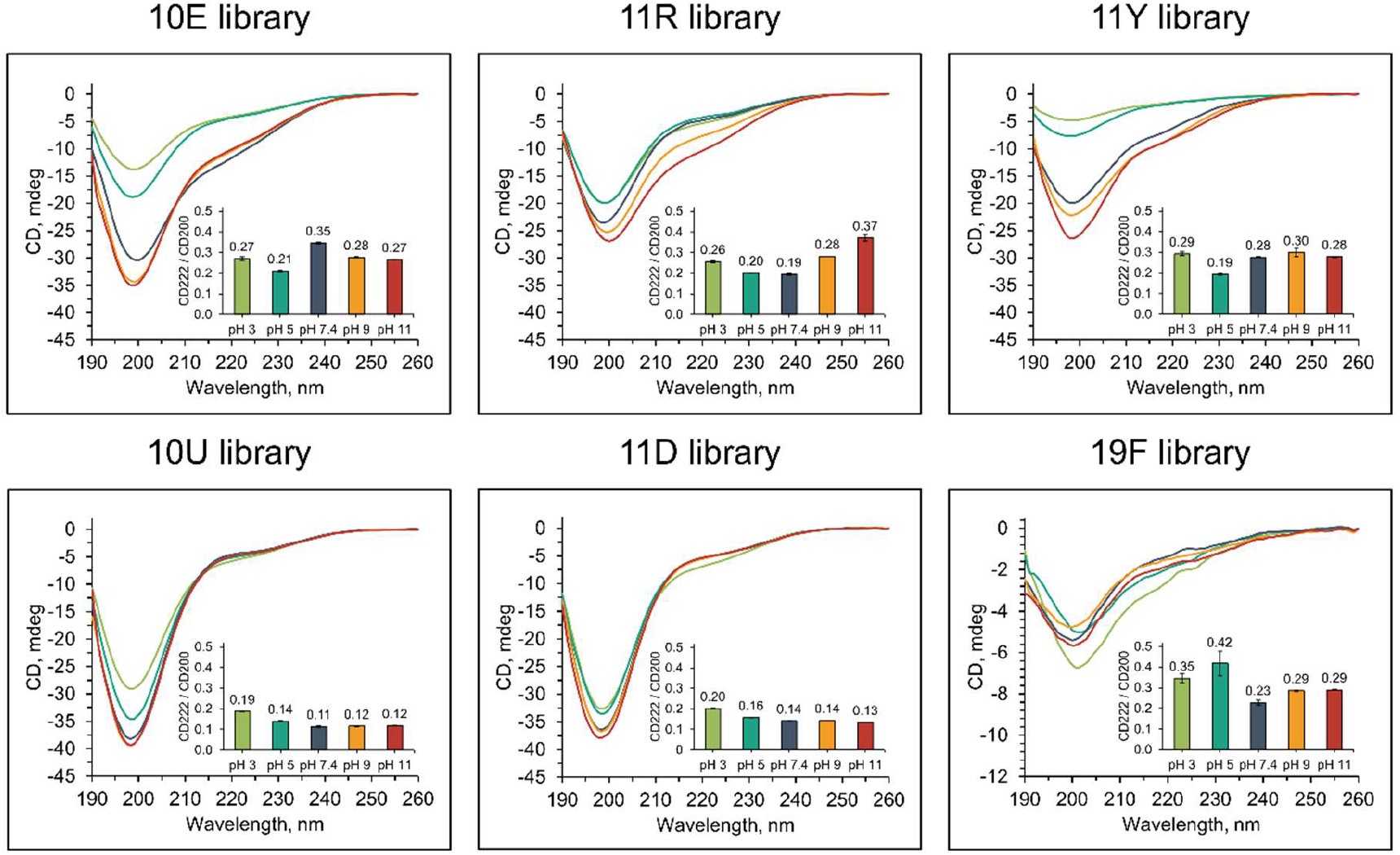
The effect of pH on the secondary structure of 25-mer combinatorial peptide libraries. CD spectra were recorded at a nominal concentration of 0.2 mg/ml peptide library (however is less for aggregation prone libraries, e.g. 19F) in a series of 10 mM ABP buffers (pH 3–11) at 25 mM NaCl. The inlet graph shows the ratios of CD signals at 222 nm to 200 nm.

**Figure 4.**
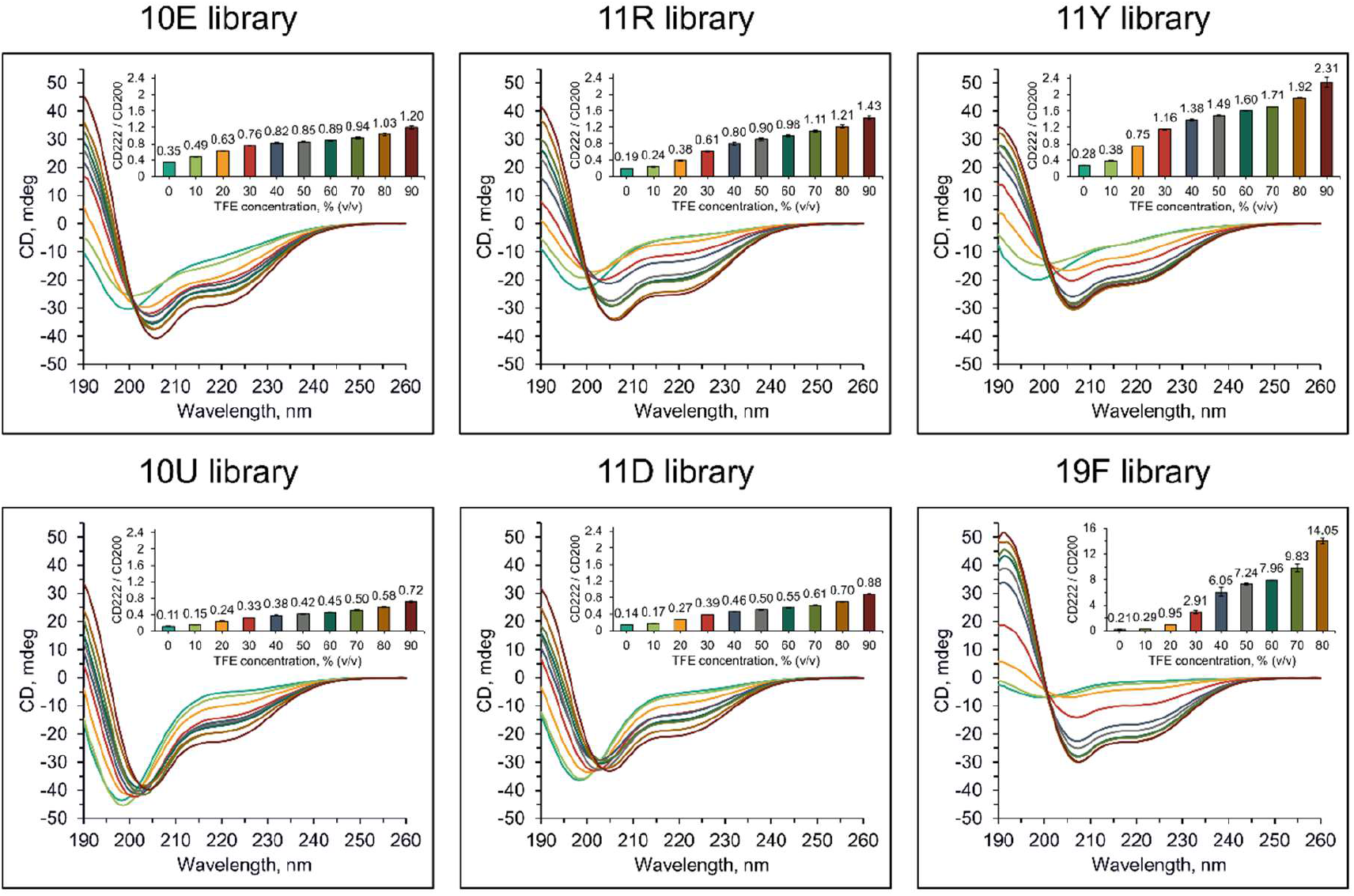
The effect of 2,2,2-trifluoroethanol to induce secondary structure of 25-mer combinatorial peptide libraries. CD spectra were recorded at 0.2 mg/ml nominal concentration in 10 mM ABP buffer (pH 7.4) with 0–90% (v/v) 2,2,2-trifluoroethanol. The inlet graphs show the ratio of CD signal at 222 to 200 nm. The y-axis has the same scale for all except for 19F library.

The CD spectra of the 10U and 11D libraries showed significantly lower relative signals at 222 nm, indicating that these peptides possessed profoundly decreased secondary structural propensity when compared with the respective canonical libraries 10E and 11R (Figure 3). The 10E library with branched aliphatic amino acids has a higher structural propensity than the 10U library with unbranched aliphatic alternatives across the whole pH range. The addition of DAB in the 11D library also significantly decreases secondary structure propensity while the addition of canonical amino acids, Arg (in 11R) or Tyr (in 11Y), increases the secondary structure propensity mildly under some pH values. This trend becomes more pronounced upon TFE titration which is often used to induce helical structure.^27–29^ TFE addition potentiates secondary structure content robustly in the 10E library, an effect that is further amplified in alphabets that include Arg and Tyr (Figure 4). Remarkably, the inclusion of DAB (in the 11D library) significantly decreases the potentiating capacity of TFE, implying that the inclusion of DAB reduces the secondary structure potential. While the poor solubility of the 19F library is reflected in the overall low intensity of its CD spectra, its elevated structural propensity (when compared with all the other library subsets) becomes evident in the TFE titration (Figure 4).

## DISCUSSION

The absence of ABA, Nva, and Nle in the proteinogenic alphabet has been considered striking given their high prebiotic abundance.^30^ Some authors have commented that this universal feature of biochemistry is a potential example of a “frozen accident” due to the absence of an obvious advantage of the canonical aliphatic residues over these ncAAs. At the same time, Weber and Miller hypothesized that inclusion of linear aliphatic amino acids would increase the side chain mobility and hence would not promote formation of ordered tertiary structure.^9^ Our experimental observations support this hypothesis. Moreover, structural prediction of 1,000 randomly chosen peptides by PEPstrMOD (using molecular dynamics based on force field libraries) confirms the same trend (Figure 5).^31^

**Figure 5.**
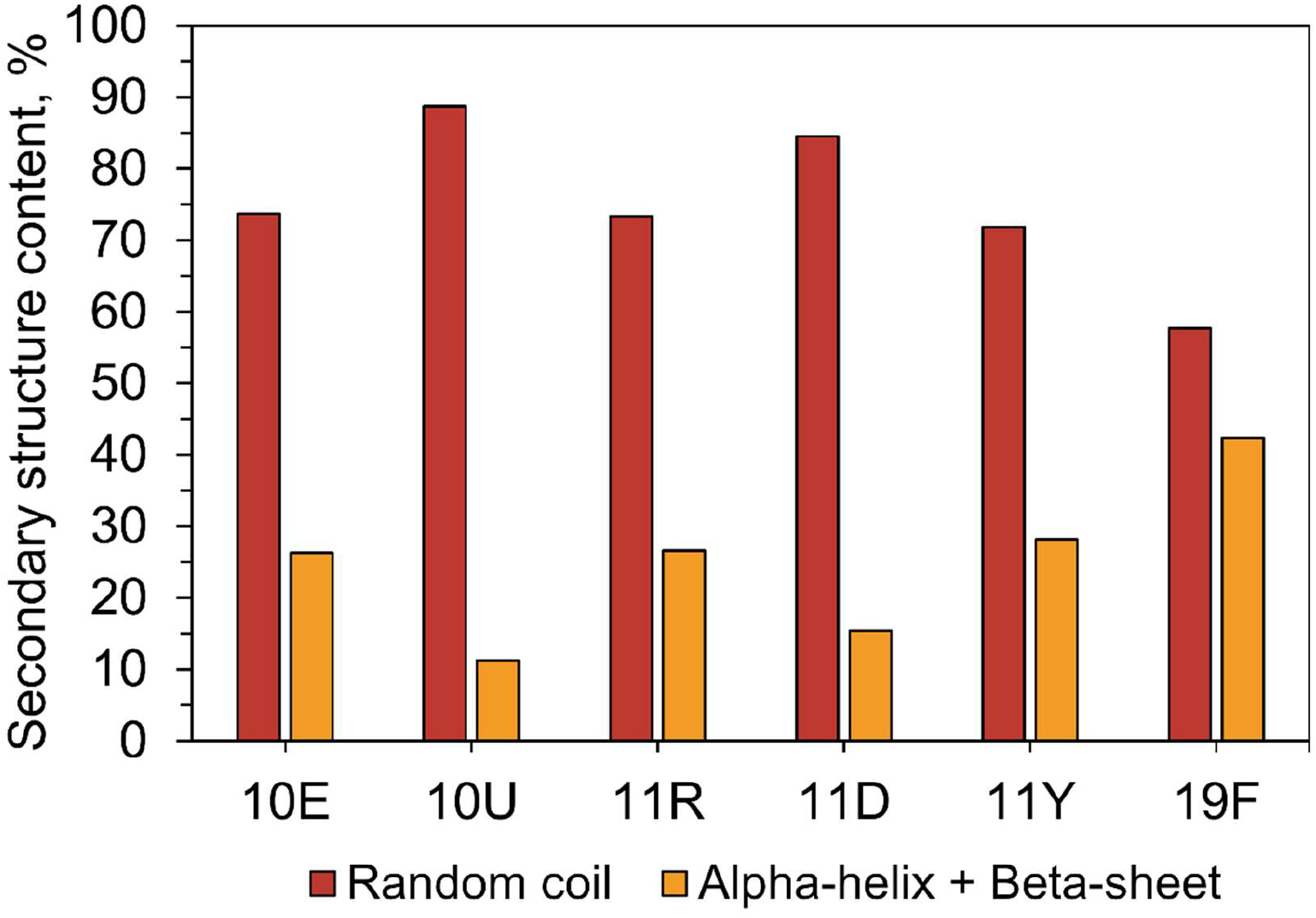
PEPstrMOD prediction of secondary structure content of 1,000 sequences selected randomly from the 25-mer combinatorial peptide libraries.

Even though unbranched amino acids are slightly more hydrophobic than their branched peers, they are expected to be much less adept at packing internal hydrophobic cores, a prerequisite to fold globular domains.^32,33^ The model shown in Figure 6A neatly explains why the 10U library has lower secondary structure-forming potential. Specifically, unbranched hydrocarbons have more conformational freedom than their branched isomers, therefore the ordering that is implicit in packing is expected to be more entropically costly. For small peptides, the hydrogen bonding associated with secondary structure is normally isoergonic between unfolded and folded states, so the driving force for secondary structure formation arises from coupling to hydrophobic collapse.^32,34–36^ The lower packing efficiency of these side chains simultaneously explains the high solubility of the 10U library. Hence, the requirement of foldability most likely acted to purge ABA, Nva, and Nle from the pool of amino acids that ultimately became canonical.

**Figure 6.**
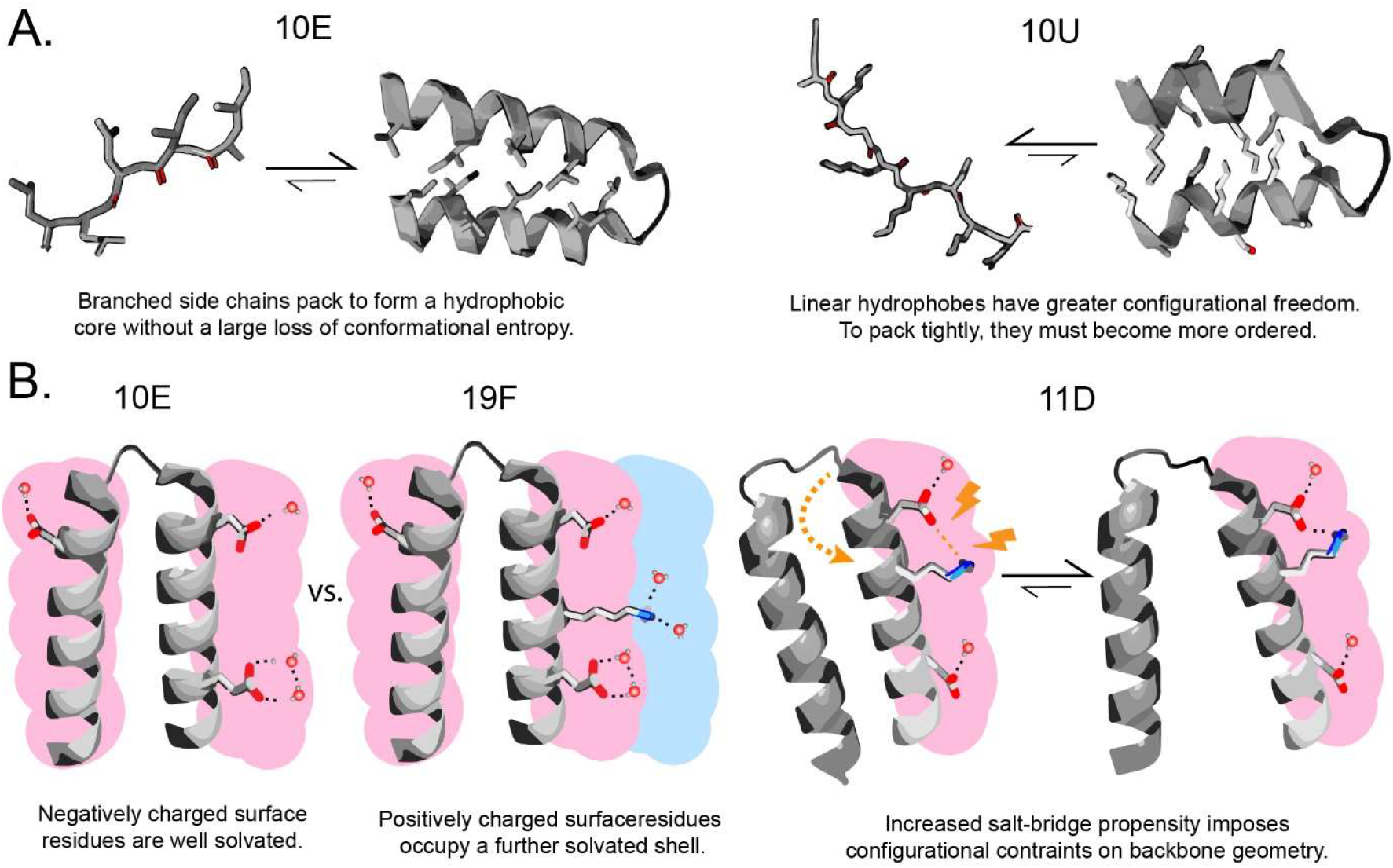
Biophysical models for the detrimental effect of unbranched hydrophobic sidechains (A) and short-chained basic residues (B) on folding stability of small proteins. (A) Unbranched hydrophobic residues render core packing more entropically expensive. (B) Short-chained basic residues occupy the same solvation sphere as canonical acidic residues, increasing the frequency of salt bridges over solvation with water.

The more prebiotically abundant diamino acids, DAP and DAB, are also strikingly absent in the modern amino acid alphabet. One possible scenario is that DAP and DAB were utilized in an ancient alphabet, and then were ultimately displaced with Arg, Lys, and His.^11^ Another school of thought holds that proteins lacked positively charged residues altogether until biosynthetic pathways for the prebiotically-unavailable Arg, Lys and His evolved.^8,37^ While the second theory might seem unlikely at first glance, proteins lacking positively charged amino acids have recently been shown capable of folding and even binding with nucleic acids, assisted by metal ion cofactors.^22,38–40^ Our results concerning the 11D library further support the second theory. Hence, this body of work is building a compelling case that primordial proteins were highly acidic.

We find that when DAB is incorporated into the 10E alphabet, it reduces secondary structure potential. Hence, its inclusion would have been a step backward rather than a step forward toward the goal of assembling foldable polypeptides. The same trend is observed in the PEPstrMOD predictions (Figure 5). The 11D library is also exceptionally soluble, so the combination of high solubility and low foldability implies that 11D polypeptides are mostly unfolded in aqueous solution.

Why does DAB harm protein structure propensity so significantly? It has been previously pointed out that these short-chain amines suffer from lactamization and acyl migration in peptides.^9,41^ This point is important in relation to the long-term stability of peptide chains, but it cannot feasibly explain the results of our study given that lactamization is a slow reaction in relation to the timescale of our experimental work. Moreover, the MALDI spectra of the library did not show any detectable scission products. A second possibility that we propose (Figure 6B) is that DAB disrupts structural potential because of a conflict associated with its simultaneous presence with Asp and Glu, i.e., negatively charged AAs with short sidechains. Especially in these short 25-mer polypeptides, charged residues are likely to be on the surface. However, by having surface-anions and surface-cations in such close proximity, a polypeptide would be inclined to fold in such a way to favour formation of ion-pair salt bridges (Figure 6B, right). The backbone angles required to form such salt bridges would create additional constraints that might be hard to simultaneously satisfy along with hydrophobic packing, hence the ion-pairs would stabilize the protein in an unpacked (and therefore unfolded) conformation. Naturally, there are two ways to avoid this folding problem: Either (1) place positive charges further away from the backbone (on a longer sidechain) so that they will primarily interact by hydrogen bonding with water in a distinct solvation shell; or (2) do not include positively charged residues at all (Figure 6). We hypothesize that early proteins avoided ion pair-induced unfolding using the second strategy, whereas modern proteins adopted the first, though this strategy only became available once Arg and Lys could be biosynthesized. One experiment that could further test this hypothesis is the synthesis of a counterfactual peptide library in which basic residues use short side chains (diaminobutyric acid) whilst acidic residues use long side chains (e.g., 2-aminoadipic acid). Our model would predict that such a combination would also support secondary structure formation.

In summary, our study on the solubility and secondary structure propensity of several prebiotically-relevant amino acid alphabets supports the assertion that foldability played a critical early role in governing which amino acids ultimately became part of the canonical alphabet. Unbranched amino acids and short-chain basic amino acids were excluded, despite their prebiotic abundance because they ‘over-solubilized’ polypeptides by stabilizing their unfolded conformations. More broadly, this study supports the view that the early canonical alphabet (10E), despite its deficiencies in relation to the modern canonical alphabet, was remarkably adaptive at supporting folding for the earliest proteins.

## EXPERIMENTAL PROCEDURES

All chemicals if not stated otherwise were purchased from Sigma Aldrich.

### Design and preparation of combinatorial peptide libraries

Split and mix synthesis of peptide libraries has its limitations.^42^ Complete library of 25-mers composed of 10 amino acids would in theory require 10^25^ beads. One gram of solid phase support contains about 10^6^ beads – therefore it would cover only 10^-21^%of possible structures. To increase the probability of completeness of the possible structures it is necessary to conduct the synthesis considering individual molecules.

Average molecular weight of a 25-mer peptide composed of 10 amino acids is about 2,500 Daltons. Therefore 2.5 kg of these peptides would potentially contain 6×10^23^ individual peptides. In order to have a chance to synthesize a complete library, we would need 250 kg of peptides, where each peptide would be present only once. But the chance of finding any given peptide in the mixture is about 70%. Therefore, studying a complete library is practically impossible.

We decided to synthesize the peptide mixture on 200 mg of the solid support and therefore create theoretically only 10^-4^% of possible structures. The approach using individual molecules is still capable of production of 10^17^×more possible structures than split and mix strategy. However, considering complete randomness of the synthetic process, we concluded that the sample is a representative collection of possible structures. One mg of synthetic peptide mixture contains about 2.4×10^18^ individual peptides.

Synthesis of the peptide mixture was accomplished by coupling mixtures of amino acids in which the ratio of individual amino acids was adjusted according to their reactivities. This approach was used by several authors, and various amino acid ratios were reported.^43^ We have used as the basis the ratios reported by Santi et al.^44,45^ and adjusted the ratios for unnatural amino acids based on their structure and our previous experience (Supplementary Table S1).

Synthesis was performed on the automatic peptide synthesizer Spyder Mark IV, using standard Fmoc peptide synthesis protocol.^46^ Fmoc amino acids were individually weighted (see Supplementary Table S1), mixed together and dissolved to create 0.3 M solution in 0.3 M N-hydroxybenzotriazole (HOBt) in dimethylformamide (DMF). Rink resin (200 mg, 0.42 mmol/g) was distributed into 10 ml syringes of the synthesizer and swelled in DMF for 10 minutes. The mixtures from Supplementary Table S1 were defined as extra amino acids 1 to 6 and placed into amino acid containers 21 to 26. Synthetic protocol was as follows: 2 × 1 min washing with DMF, 1 × 1 min and 1 × 20 min treatment with 20% piperidine in DMF, 4 × 1 min wash with DMF, 1 × 1 min wash with 0.3 M HOBt, 2 × 60 min coupling with amino acid mixture and 1 M diisopropylcarbodiimide in DMF. After 25 cycles of the synthesis, the resin was washed with DMF and dichloromethane and dried *in vacuo*. Dried resin was treated with 5 ml Mixture K (trifluoroacetic acid-thioanisol-water-phenol-ethanedithiol, 82.5:5:5:5:2.5, v/v) for 2 hours.^47^ Resin was filtered off and peptide mixture was precipitated with diethyl ether three times and dried *in vacuo*. The pellet was then dissolved in 20% acetic acid and lyophilized. Prior to further experiments, all the peptide libraries were lyophilized three times with 1 mM HCl overnight.

### Quality control of peptide libraries

The molecular weight distributions of combinatorial peptide libraries were confirmed by mass spectrometry using UltrafleXtreme MALDI-TOF/TOF mass spectrometer (Bruker, Germany) according to the standard procedure.

Prior to the amino acid analysis, the library samples were hydrolyzed in 6 M hydrochloric acid at 110 °C for 20 h, the hydrolysate was evaporated, and reconstituted with 0.1 M hydrochloric acid containing the internal standard. Amino acid analysis was performed on an Agilent 1260 HPLC (Agilent Technologies) equipped with a fluorescence detector using automated o-phtalaldehyde/2-mercaptopropionic acid (OPA/MPA) or ninhydrin derivatization in case of 10E, 10U, 11R and 11D, 11Y, 19F libraries, respectively.

The presence of 2,4-diaminobutyric acid in the 11D library was confirmed by fluorescamine assay.^48^ For the assay, peptide library solution (75 μl) in PBS buffer was incubated with 25 μl of fresh stock of 3 mg/ml fluorescamine in DMSO at room temperature for 1 hour in a 96-well plate (Greiner 650209). Fluorescence intensity (λEx = 365 nm, λEm = 470 nm) was recorded using Tecan Spark plate reader. Primary amine concentrations were determined by linear interpolation of measured intensities for peptide H-GTIQPYPFSWGY-NH2 in concentration range 87.5 – 2.7 μM. Obtained amine concentrations were then divided by the concentration of the analyzed peptide determined by amino acid analysis to calculate the amount of amine equivalents per molecule. Experiments were conducted in duplicate.

### Solubility and aggregation propensity measurements

A series of ten Britton-Robinson buffers at pH 3.0, 5.0, 7.4, 9.0, and 11.0 was prepared by mixing 20 mM acetic acid, 20 mM boric acid and 20 mM phosphoric acid and adjusting pH to the desired value with 5 M NaOH. The ionic strength was adjusted to either 50 mM (low ionic strength) or 500 mM (high ionic strength) with NaCl, and all Britton-Robinson buffers were filter-sterilized using 0.22 μm PVDF membrane before use. Lyophilized peptide libraries were thoroughly resuspended in autoclaved MilliQ water to 5 mg/ml and subsequently diluted ten times into Britton-Robinson buffer to the final concentration of 0.5 mg/ml. The samples were gently shaken at room temperature for 30 min and then centrifuged at maximum speed (21,300 × g) for 15 min at 4 *°C* in order to remove insoluble fraction. The supernatant (soluble fraction) was taken and used for solubility and aggregation propensity analyses.

The relative amount of peptides in soluble fractions was estimated spectrophotometrically by absorption of peptide bonds at 215 nm for all peptide libraries except the 19F peptide library. Due to the extremely low solubility, solubility of 19F peptide library was estimated by fluorescence spectroscopy using Pierce Quantitative Fluorometric Peptide Assay (Thermo Fisher Scientific, USA), and 0.5 mg/ml solution in DMSO was used as a standard. Spectrophotometric measurements were performed five times, whereas fluorometric measurements were performed in triplicates.

The relative amount of soluble aggregates was estimated by size-exclusion chromatography. The 100 μl aliquot of supernatant was loaded onto Superdex 75 Increase 10/300 GL column (Cytiva, USA) that was pre-equilibrated with two bed volumes of the corresponding 20 mM Britton-Robinson buffer. The peptides were eluted from the column by one bed volume of the buffer at 0.5 ml/min flow rate at room temperature, and the eluted peptides were detected by absorption at 215 nm.

### Circular Dichroism spectroscopy

The CD spectra were recorded using a Chirascan-plus spectrophotometer (Applied Photophysics, UK) over the wavelength range 190–260 nm in steps of 1 nm with an averaging time of 1 sec per step. Cleared samples at 0.2 mg/ml nominal concentration in 1 mm path-length quartz cells were placed into a cell holder, and spectra were recorded at room temperature. The CD signal was obtained as ellipticity in units of millidegrees and the resulting spectra were averaged from two scans and buffer-spectrum subtracted. All CD measurements were performed twice, and the resulting ratio of ellipticity at 222 nm to ellipticity at 200 nm was averaged and used to estimate the secondary structure content in peptide libraries.

To estimate the effect of pH on secondary structure, 5 mg/ml suspensions of peptide libraries in MilliQ water were diluted into 10 mM Britton-Robinson buffers at pH 3.0; 5.0; 7.4; 9.0 and 11.0 to the final concentration of 0.2 mg/ml, gently mixed at room temperature for 30 min and then centrifuged at maximum speed (21,300 × g) for 15 min at 4 *°C* in order to remove insoluble fraction.

To estimate the effect of 2,2,2-trifluoroethanol on secondary structure, 5 mg/ml suspensions of peptide libraries in MilliQ water were diluted in a series of 10 mM Britton-Robinson buffer (pH 7.4) containing 0–90% (v/v) 2,2,2-trifluoroethanol.

To estimate the effect of metal ions on secondary structure, 5 mg/ml suspensions of peptide libraries in MilliQ water were diluted in a series of 10 mM Tris buffer at pH 7.4 supplemented with 0; 10; 100; and 1,000 μM mixture of NaCl, KCl, MgCl2, MnCl2, and ZnCl2.

### Bioinformatic analysis of in silico peptide libraries

1,000 25-mer sequences were randomly generated *in silico* for the libraries and their structure was predicted using PEPstrMOD.^31^ PEPstrMOD uses force field libraries Forcefield_NCAA and Forcefield_PTM and integrates them in Molecular Dynamics (MD) simulations using AMBER. Additionally, it implements the force field library SwissSideChain with the MD package GROMACS. The parameters employed in the MD simulations were 50 picoseconds of simulation time and the selected peptide environment was set to “vacuum”.

DSSP^49^ was employed to determine the secondary structure elements from the peptide tertiary structures generated by PEPstrMOD. The obtained annotations of secondary structure were grouped into three larger classes: helix (H: alpha-helix, G: 310 helix and I: π-helix), sheet (B: isolated β-bridge and E: extended strand) and loop (T: hydrogen bonded turn, S: bend and “_” blank spaces).

## ASSOCIATED CONTENT

### Supporting Information

Supplementary Table S1 and Supplementary Figures S1-S6.

## AUTHOR INFORMATION

### Corresponding Authors

**Stephen D. Fried** *Department of Chemistry, Johns Hopkins University, Baltimore, Maryland 21218, United States; T. C. Jenkins Department of Biophysics, Johns Hopkins University, Baltimore, Maryland 21218, United States*

**Klara Hlouchova** *Department of Cell Biology, Faculty of Science, Charles University, BIOCEV, Prague, 12843, Czech Republic; Institute of Organic Chemistry and Biochemistry, Czech Academy of Sciences, Prague, 16610, Czech Republic*

### Authors

**Mikhail Makarov** *Department of Cell Biology, Faculty of Science, Charles University, BIOCEV, Prague, 12843, Czech Republic*

**Alma C. Sanchez Rocha** *Department of Cell Biology, Faculty of Science, Charles University, BIOCEV, Prague, 12843, Czech Republic*

**Robin Krystufek** *Department of Physical Chemistry, Faculty of Science, Charles University, Prague, 12843, Czech Republic; Institute of Organic Chemistry and Biochemistry, Czech Academy of Sciences, Prague, 16610, Czech Republic*

**Ivan Cherepashuk** *Department of Cell Biology, Faculty of Science, Charles University, BIOCEV, Prague, 12843, Czech Republic*

**Volha Dzmitruk** *Institute of Biotechnology of the Czech Academy of Sciences, BIOCEV, Vestec, 25250, Czech Republic*

**Tatsiana Charnavets** *Institute of Biotechnology of the Czech Academy of Sciences, BIOCEV, Vestec, 25250, Czech Republic*

**Anneliese M. Faustino** *Department of Chemistry, Johns Hopkins University, Baltimore, Maryland 21218, United States*

**Michal Lebl** *Institute of Organic Chemistry and Biochemistry, Czech Academy of Sciences, Prague, 16610, Czech Republic*

**Kosuke Fujishima** *Earth-Life Science Institute, Tokyo Institute of Technology, Tokyo, 1528550, Japan; Graduate School of Media and Governance, Keio University, Fujisawa, 2520882, Japan*

### Notes

The authors declare no competing interests.

## ACKNOWLEDGEMENTS

We would like to thank Dr. Vyacheslav Tretyachenko and Dr. Valerio Guido Giacobelli for helpful discussions about this project. We would also like to acknowledge Dr. Radko Souček and Dr. Martin Hubálek for their assistance with amino acid analysis and MALDI analyses, respectively. This work was supported by the Human Frontier Science Program grant HFSP-RGY0074/2019 to K.F., S.D.F., and K.H. M.M., A.C.S.R., and I.C. acknowledge support by the project the “Grant Schemes at CU” (reg. no. CZ.02.2.69/0.0/0.0/19_073/0016935), project no. START/SCI/148. I.C. is supported by the Visegrad Fund Scholarship (no. 52110039). V.D., T.Ch., K.H., M.M., A.C.S.R., and I.C. acknowledge CMS CF Biophysical techniques of CIISB, Instruct-CZ Centre, supported by MEYS CR Infrastructure project LM2018127 and European Regional Development Fund-Project „UP CIISB‟ (No. CZ.02.1.01/0.0/0.0/18_046/0015974). S.D.F. acknowledges support from the NIH Director’s New Innovator Award (DP2-GM140926).

## REFERENCES

1. Milner-White, E. J.; Russell, M. J. Functional Capabilities of the Earliest Peptides and the Emergence of Life. Genes (Basel). 2011, 2 (4), 671–688. DOI: 10.3390/genes2040671.

2. Fried, S. D.; Fujishima, K.; Makarov, M.; Cherepashuk, I.; Hlouchova, K. Peptides Before and During the Nucleotide World: An Origins Story Emphasizing Cooperation Between Proteins and Nucleic Acids. J. R. Soc. Interface. 2022, 19 (187), 20210641. DOI: 10.1098/rsif.2021.0641

3. Frenkel-Pinter, M.; Frenkel-Pinter, M.; Samanta, M.; Ashkenasy, G.; Leman, L. J. Prebiotic Peptides: Molecular Hubs in the Origin of Life. Chem. Rev. 2020, 120 (11), 4707–4765. DOI: 10.1021/acs.chemrev.9b00664

4. Freeland, S. Undefining Life’s Biochemistry: Implications for Abiogenesis. J. R. Soc. Interface. 2022, 19 (187), 20210814. DOI: 10.1098/rsif.2021.0814

5. Cleaves, H. J. The Origin of the Biologically Coded Amino Acids. J. Theor. Biol. 2010, 263 (4), 490–498. DOI: 10.1016/j.jtbi.2009.12.014

6. Kitadai, N.; Maruyama, S. Origins of building blocks of life: A review. Geosci. Front. 2018, 9 (4), 1117–1153. DOI: 10.1016/j.gsf.2017.07.007

7. Trifonov, E. N. The Triplet Code from First Principles. J. Biomol. Struct. Dyn. 2004, 22 (1), 1–11. DOI: 10.1080/07391102.2004.10506975

8. Higgs, P. G.; Pudritz, R. E. A Thermodynamic Basis for Prebiotic Amino Acid Synthesis and the Nature of the First Genetic Code. Astrobiology. 2009, 9 (5), 483–490. DOI: 10.1089/ast.2008.0280

9. Weber, A. L.; Miller, S. L. Reasons for the Occurrence of the Twenty Coded Protein Amino Acids. J. Mol. Evol. 1981, 17 (5), 273–284. DOI: 10.1007/BF01795749

10. Freeland, S. “Terrestrial” Amino Acids and Their Evolution. In Amino Acids, Peptides and Proteins in Organic Chemistry: Origins and Synthesis of Amino Acids, Volume 1, 1st ed.; Wiley-VCH Verlag GmbH & Co. KGaA, 2010; pp 43–75.

11. Vázquez-Salazar, A.; Lazcano, A. Early Life: Embracing the RNA World. Curr. Biol. 2018, 28 (5):R220–222. DOI: 10.1016/j.cub.2018.01.055

12. Raggi L.; Bada, J. L.; Lazcano, A. On the Lack of Evolutionary Continuity between Prebiotic Peptides and Extant Enzymes. Phys. Chem. Chem. Phys. 2016, 18 (30), 20028–20032. DOI: 10.1039/C6CP00793G

13. Bredehöft, J.; Thiemann, W.; Jessberger, E.; Carob, G.; Meierhenrich, U. Identification of Diamino Acids in the Murichison Meteorite. Proc. Natl. Acad. Sci. U. S. A. 2004, 101 (25), 9182–9186. DOI: 10.1073/pnas.0403043101

14. Wolman, Y.; Haverland, W. J.; Miller, S. L. Nonprotein Amino Acids from Spark Discharges and Their Comparison with the Murchison Meteorite Amino Acids. Proc. Natl. Acad. Sci. U. S. A. 1972, 69 (4), 809–811. DOI: 10.1073/pnas.69.4.809

15. Alvarez-Carreño, C.; Becerra, A.; Lazcano, A. Norvaline and Norleucine May Have Been More Abundant Protein Components during Early Stages of Cell Evolution. Orig. Life Evol. Biosph. 2013, 43 (4–5), 363–375. DOI: 10.1007/s11084-013-9344-3

16. Apostol, I.; Levine, J.; Lippincott, J.; Leach, J.; Hess, E.; Glascock, C. B.; Weickert, M. J.; Blackmore, R. Incorporation of Norvaline at Leucine Positions in Recombinant Human Hemoglobin Expressed in Escherichia Coli. J. Biol. Chem. 1997, 272 (46), 28980–28988. DOI: 10.1074/jbc.272.46.28980

17. Martin, W.; Russell M. J. On the Origin of Biochemistry at an Alkaline Hydrothermal Vent. Philos. Trans. R. Soc. Lond. B. Biol. Sci. 2007, 362 (1486), 1887–1925. DOI: 10.1098/rstb.2006.1881

18. Grotzinger, J. P.; Kasting, J. F. New Constraints on Precambrian Ocean Composition. J. Geol. 1993, 101 (2), 235–243. DOI: 10.1086/648218

19. Tanaka, J.; Doi, N.; Takashima, H.; Yanagawa, H. Comparative Characterization of Random-Sequence Proteins Consisting of 5, 12, and 20 Kinds of Amino Acids. Protein Sci. 2010, 19 (4), 786–795. DOI: 10.1002/pro.358

20. Newton, M. S.; Morrone, D. J.; Lee, K. H.; Seelig, B. Genetic Code Evolution Investigated through the Synthesis and Characterisation of Proteins from Reduced-Alphabet Libraries. ChemBioChem. 2019, 20 (6), 846–856. DOI: 10.1002/cbic.201800668

21. Tretyachenko, V.; Vymětal, J.; Bednárová, L.; Kopecký, V.; Hofbauerová, K.; Jindrová, H.; Hubálek, M.; Souček, R.; Konvalinka, J.; Vondrášek, J.; Hlouchová, K. Random Protein Sequences Can Form Defined Secondary Structures and Are Well-Tolerated In Vivo. Sci. Rep. 2017, 7 (1), 15449. DOI: 10.1038/s41598-017-15635-8

22. Tretyachenko, V.; Vymětal, J.; Neuwirthová, T.; Vondrášek, J.; Fujishima, K.; Hlouchová, K. Modern and Prebiotic Amino Acids Support Distinct Structural Profiles in Proteins. Open Biol. 2022, 12 (6), 220040. DOI: 10.1098/rsob.220040

23. Kozlowski, L. P. Proteome-pI: Proteome Isoelectric Point Database. Nucleic Acids Res. 2017, 45 (D1), D1112–D1116. DOI: 10.1093/nar/gkw978

24. Wang, Y.; Van Oosterwijk, N.; Ali, A. M.; Adawy, A.; Anindya, A. L.; Dömling, A. S. S.; Groves M. R. A Systematic Protein Refolding Screen Method using the DGR Approach Reveals that Time and Secondary TSA are Essential Variables. Sci. Rep. 2017, 7 (1), 9355. DOI: 10.1038/s41598-017-09687-z

25. Reynolds, J. A.; Gilbert, D. B.; Tanford, C. Empirical Correlation Between Hydrophobic Free Energy and Aqueous Cavity Surface Area. Proc. Natl. Acad. Sci. U. S. A. 1974, 71 (8), 2925–2927. DOI: 10.1073/pnas.71.8.2925

26. Vymětal, J.; Vondrášek, J.; Hlouchová, K. Sequence Versus Composition: What Prescribes IDP Biophysical Properties? Entropy (Basel). 2019, 21 (7), 654. DOI: 10.3390/e21070654

27. Kelly, S. M.; Jess, T. J.; Price, N. C. How to Study Proteins by Circular Dichroism. Biochim. Biophys. Acta. 2005, 1751 (2),119–139. DOI: 10.1016/j.bbapap.2005.06.005

28. Buck, M. Trifluoroethanol and Colleagues: Cosolvents Come of Age. Recent Studies with Peptides and Proteins. Q. Rev. Biophys. 1998, 31 (3), 297–355. DOI: 10.1017/s003358359800345x

29. Starzyk, A.; Barber-Armstrong, W.; Sridharan, M.; Decatur, S. M. Spectroscopic Evidence for Backbone Desolvation of Helical Peptides by 2,2,2-Trifluoroethanol: An Isotope-Edited FTIR Study. Biochemistry. 2005, 44 (1),369–376. DOI: 10.1021/bi0481444

30. Zaia, D. A. M.; Zaia, C. T. B. V.; De Santana H. Which Amino Acids Should Be Used in Prebiotic Chemistry Studies? Orig. Life Evol. Biosph. 2008, 38 (6), 469–488. DOI: 10.1007/s11084-008-9150-5

31. Singh, S.; Singh, H.; Tuknait, A.; Chaudhary, K.; Singh, B.; Kumaran, S.; Raghava, G. P. PEPstrMOD: Structure Prediction of Peptides Containing Natural, Non-Natural and Modified Residues. Biol. Direct. 2015, 10 (1), 73. DOI: 10.1186/s13062-015-0103-4

32. Tanford, C. The Hydrophobic Effect and The Organization of Living Matter. Science. 1978, 200 (4345), 1012–1018. DOI: 10.1126/science.653353

33. Dill, K. A. Dominant Forces in Protein Folding. Biochemistry. 1990, 29 (31), 7133–7155. DOI: 10.1021/bi00483a001

34. Klotz, I. M.; Franzen, J. S. Hydrogen Bonds between Model Peptide Groups in Solution. J. Am. Chem. Soc. 1962, 84 (18), 3461–3466. DOI: 10.1021/ja00877a009

35. Koh, J. T.; Cornish, V. W.; Schultz, P. G. An Experimental Approach to Evaluating The Role of Backbone Interactions in Proteins using Unnatural Amino Acid Mutagenesis. Biochemistry. 1997, 36 (38), 11314–11322. DOI: 10.1021/bi9707685

36. Baldwin, R. L. In Search of The Energetic Role of Peptide Hydrogen Bonds. J. Biol. Chem. 2003, 278 (20), 17581–17588. DOI: 10.1074/jbc.X200009200

37. Trifonov, E. N. Consensus Temporal Order of Amino Acids and Evolution of The Triplet Code. Gene. 2000, 261 (1), 139–151. DOI: 10.1016/s0378-1119(00)00476-5

38. Longo, L. M.; Blaber, M. Protein Design at The Interface of The Pre-Biotic and Biotic Worlds. Arch. Biochem. Biophys. 2012, 526 (1), 16–21. DOI: 10.1016/j.abb.2012.06.009

39. Despotović, D.; Longo, L. M.; Aharon, E.; Kahana, A.; Scherf, T.; Gruic-Sovulj, I.; Tawfik, D. S. Polyamines Mediate Folding of Primordial Hyperacidic Helical Proteins. Biochemistry. 2020, 59 (46), 4456–4462. DOI: 10.1021/acs.biochem.0c00800

40. Giacobelli, V. G.; Fujishima, K.; Lepšík, M.; Tretyachenko, V.; Kadavá, T.; Makarov, M.; Bednárová, L.; Novák, P.; Hlouchová K. In Vitro Evolution Reveals Noncationic Protein-RNA Interaction Mediated by Metal Ions. Mol. Biol. Evol. 2022, 39 (3), msac032. DOI: 10.1093/molbev/msac032

41. Poduska, K.; Katrukha, G. S.; Silaev, A. B.; Rudinger, J. Amino Acids and Peptides. LII. Intramolecular Aminolysis of Amide Bonds in Derivatives of Alpha,Gamma-Diaminobutyric Acid, Alpha,Beta-Diaminoproprionic Acid, and Ornithine. Collect. Czechoslov. Chem. Commun. 1965, 30, 2410–2433. DOI: 10.1135/cccc19652410

42. Lam, K. S.; Lebl, M.; Krchňák, V. The “One-Bead-One-Compound” Combinatorial Library Method. Chem. Rev. 1997, 97 (2),411–448. DOI: 10.1021/cr9600114

43. Ostresh, J. M.; Winkle, J. H.; Hamashin, V. T.; Houghten R. A. Peptide Libraries: Determination of Relative Reaction Rates of Protected Amino Acids in Competitive Couplings. Biopolymers. 1994, 34 (12), 1681–1689. DOI: 10.1002/bip.360341212

44. Rutter, W.; Santi, D. Peptide Mixtures. US 5420246 A, 1995.

45. Ivanetich, K. M.; Santi, D. V. Preparation of Equimolar Mixtures of Peptides by Adjustment of Activated Amino Acid Concentrations. Methods Enzymol. 1996, 267, 247–260. DOI: 10.1016/S0076-6879(96)67017-7

46. Lebl, M.; Flegelova, Z.; Poncar, P.; Mudra, P.; Paces, O.; Knor, M.; Buzek, M.; Smrz, J.; Pokorny, V.; Pesek, V. Device for Parallel Oligomer Synthesis, Method of Parallel Oligomer Synthesis and Use Thereof. WO-2019115514-A1, 2017.

47. King, D. S.; Fields, C. G.; Fields, G. B. A Cleavage Method Which Minimizes Side Reactions Following Fmoc Solid Phase Peptide Synthesis. Int. J. Pept. Protein Res. 1990, 36 (3), 255–266. DOI: 10.1111/j.1399-3011.1990.tb00976.x

48. Udenfriend, S.; Stein, S.; Böhlen, P.; Dairman, W.; Leimgruber, W.; Weigele, M. Fluorescamine: a Reagent for Assay of Amino Acids, Peptides, Proteins, and Primary Amines in The Picomole Range. Science. 1972, 178 (4063), 871–872. DOI: 10.1126/science.178.4063.871

49. Kabsch, W.; Sander, C. Dictionary of Protein Secondary Structure: Pattern Recognition of Hydrogen-Bonded and Geometrical Features. Biopolymers. 1983, 22 (12), 2577–2637. DOI: 10.1002/bip.360221211

