## Supplementary material for "Early selection of the amino acid alphabet was adaptively shaped by biophysical constraints of foldability"

*Supplementary Table S1: Weights (mg) and amounts (mmol) of protected amino acids in individual isokinetic mixtures used for synthesis. Mixtures were dissolved in a final volume listed in the last row yielding 0.3 M solution of amino acids in 0.3 M HOBt in DMF.*

| | 19F | | 10E | | 11R | | 11Y | | 10U | | 11D | |
|---|---|---|---|---|---|---|---|---|---|---|---|---|
| | m mg | n mmol | m mg | n mmol | m mg | n mmol | m mg | n mmol | m mg | n mmol | m mg | n mmol |
| Fmoc-Ala | 202 | 0.61 | 332 | 1.01 | 300 | 0.91 | 312 | 0.95 | 508 | 1.54 | 303 | 0.92 |
| Fmoc-Asp(OtBu) | 276 | 0.67 | 455 | 1.11 | 409 | 0.99 | 426 | 1.04 | 694 | 1.69 | 414 | 1.01 |
| Fmoc-Glu(OtBu) | 296 | 0.67 | 489 | 1.10 | 440 | 0.99 | 457 | 1.03 | 745 | 1.68 | 444 | 1.00 |
| Fmoc-Phe | 187 | 0.48 | | | | | | | | | | |
| Fmoc-Gly | 164 | 0.55 | 270 | 0.91 | 243 | 0.82 | 253 | 0.85 | 412 | 1.39 | 246 | 0.83 |
| Fmoc-His(Trt) | 422 | 0.68 | | | | | | | | | | |
| Fmoc-Ile | 1,171 | 3.31 | 1,934 | 5.47 | 1,740 | 4.92 | 1,809 | 5.12 | | | 1,759 | 4.98 |
| Fmoc-Lys(Boc) | 556 | 1.19 | | | | | | | | | | |
| Fmoc-Leu | 335 | 0.95 | 553 | 1.56 | 497 | 1.41 | 517 | 1.46 | | | 503 | 1.42 |
| Fmoc-Met | 163 | 0.44 | | | | | | | | | | |
| Fmoc-Asn(Trt) | 609 | 1.02 | | | | | | | | | | |
| Fmoc-Pro | 278 | 0.78 | 460 | 1.29 | 414 | 1.16 | 430 | 1.21 | 702 | 1.98 | 418 | 1.18 |
| Fmoc-Gln(Trt) | 620 | 1.02 | | | | | | | | | | |
| Fmoc-Arg(Pmc) | 807 | 1.22 | | | 1,199 | 1.81 | | | | | | |
| Fmoc-Ser(tBu) | 203 | 0.53 | 336 | 0.88 | 302 | 0.79 | 314 | 0.82 | 512 | 1.34 | 306 | 0.80 |
| Fmoc-Thr(tBu) | 363 | 0.91 | 599 | 1.51 | 539 | 1.36 | 560 | 1.41 | 914 | 2.30 | 545 | 1.37 |
| Fmoc-Val | 731 | 2.15 | 1,208 | 3.56 | 1,086 | 3.20 | 1,130 | 3.33 | | | 1,099 | 3.24 |
| Fmoc-Trp(Boc) | 254 | 0.48 | | | | | | | | | | |
| Fmoc-Tyr(tBu) | 363 | 0.79 | | | | | 560 | 1.22 | | | | |
| Fmoc-Nva | | | | | | | | | 752 | 2.22 | | |
| Fmoc-Nle | | | | | | | | | 775 | 2.19 | | |
| Fmoc-Aba | | | | | | | | | 648 | 1.99 | | |
| Fmoc-Dab(Boc) | | | | | | | | | | | 734 | 1.67 |
| V / ml | 61.5 | | 61.3 | | 61.2 | | 61.4 | | 39.7 | | 55.8 | |

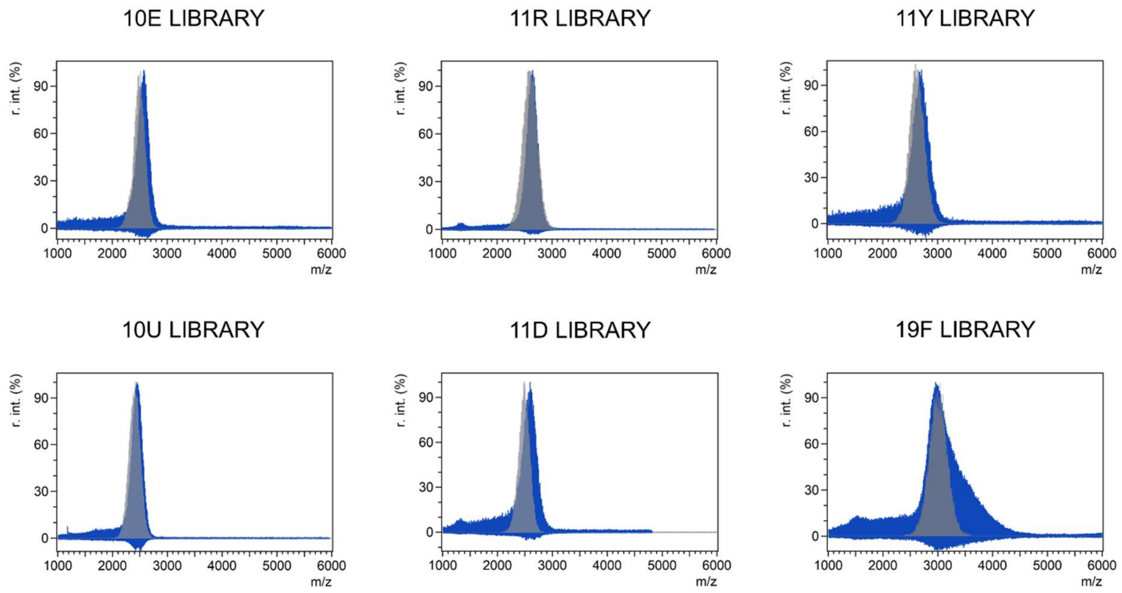

*Supplementary Figure S1. Molecular weight distribution of 25-mer combinatorial peptide libraries estimated by MALDI-TOF-MS using 2,5-dihydrobenzoic acid (DHB) matrix. Observed m/z distribution is shown in blue, expected m/z distribution is shown as a gray overlay. Peptide libraries were dissolved in acetonitrile:water (1:1) mixture + 0.1 % (v/v) acetic acid. 19F peptide library was dissolved in acetonitrile:water (1:1) mixture.*

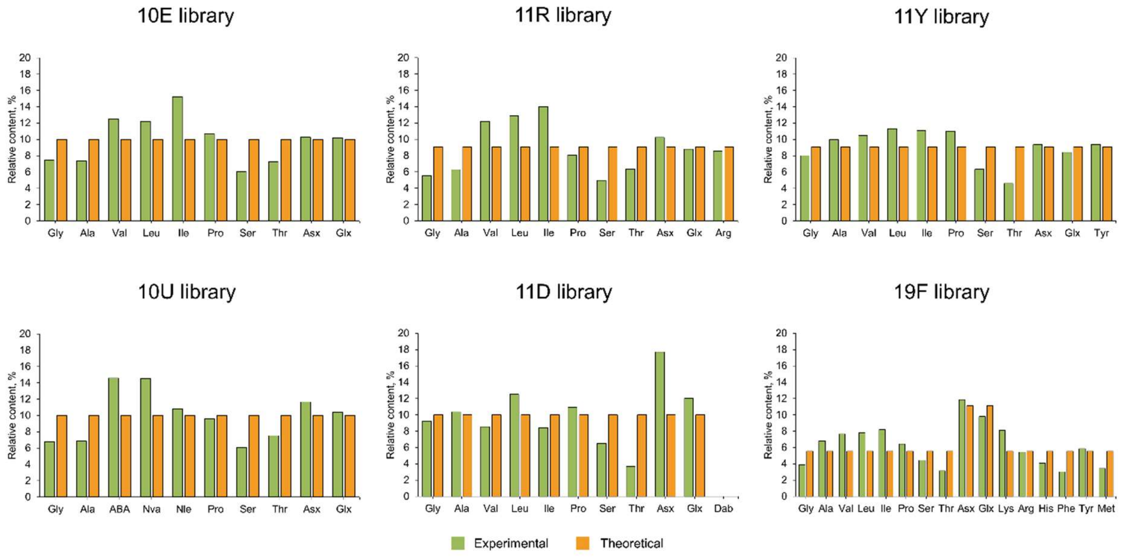

*Supplementary Figure S2. Amino acid composition of 25-mer combinatorial peptide libraries determined by amino acid analysis.*

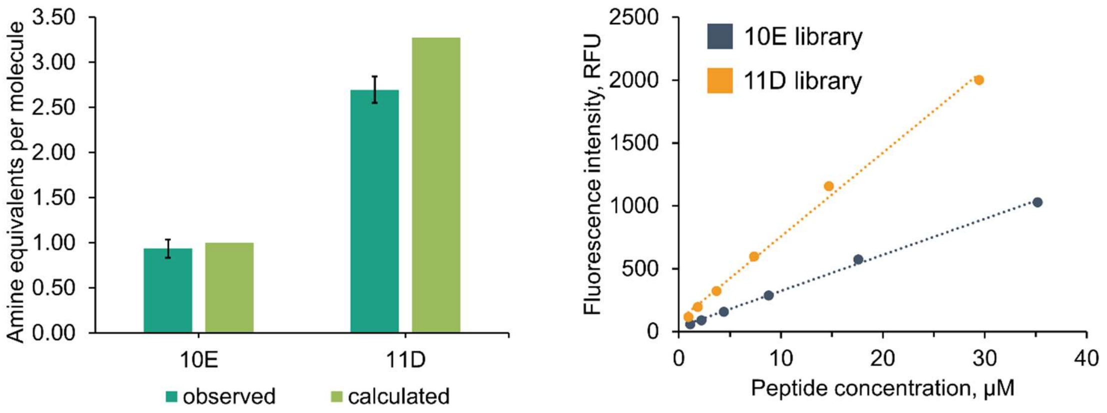

*Supplementary Figure S3. The quantification of free amino acid groups in 10E and 11D peptide libraries using fluorescamine assay.*

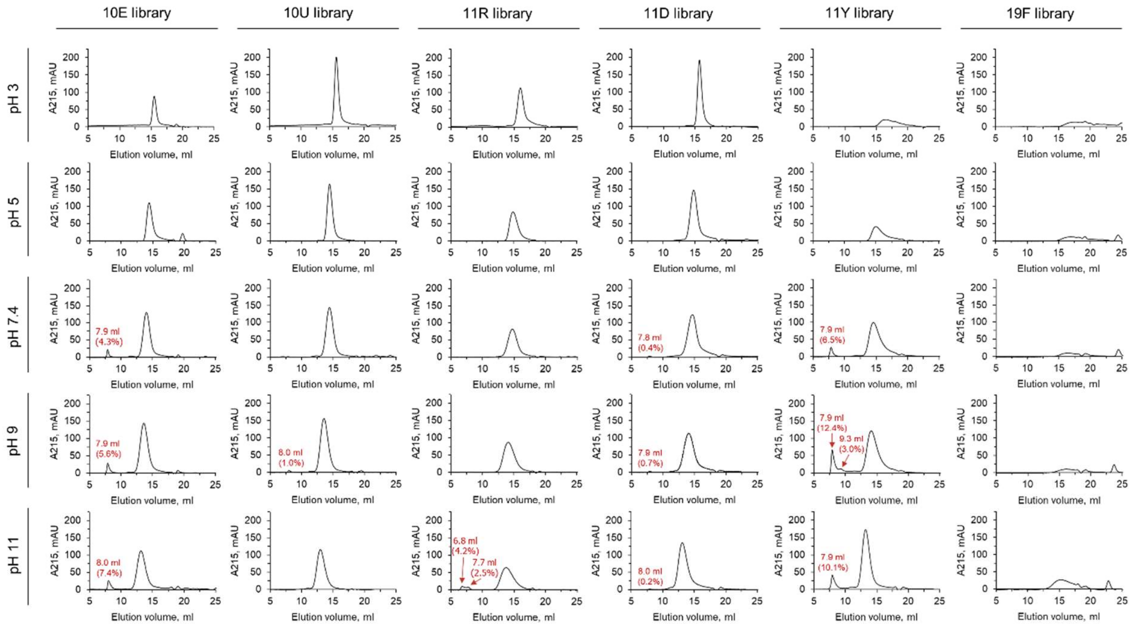

*Supplementary Figure S4. Aggregation propensity of 25-mer combinatorial peptide libraries at different pH and low ionic strength (50 mM NaCl). Aggregation propensity was measured for 0.5 mg/ml nominal peptide library solutions in a series of 20 mM ABP buffers (pH 3–11) by size-exclusion chromatography.*

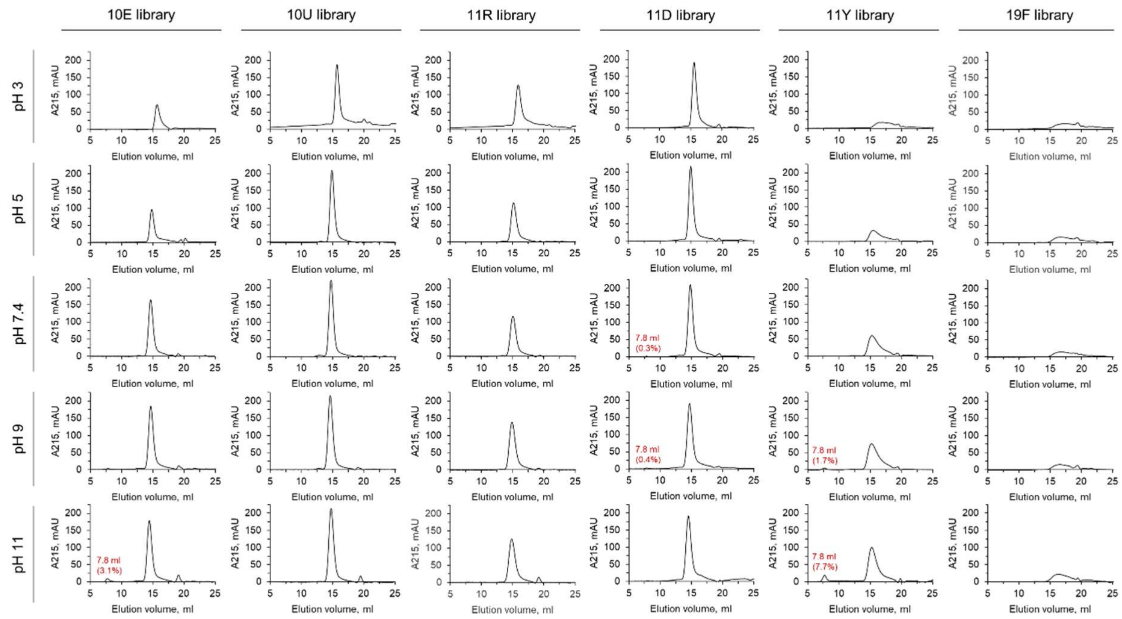

*Supplementary Figure S5. Aggregation propensity of 25-mer combinatorial peptide libraries at different pH and high ionic strength (500 mM NaCl). Aggregation propensity was measured for 0.5 mg/ml nominal peptide library solutions in a series of 20 mM ABP buffers (pH 3–11) by size-exclusion chromatography.*

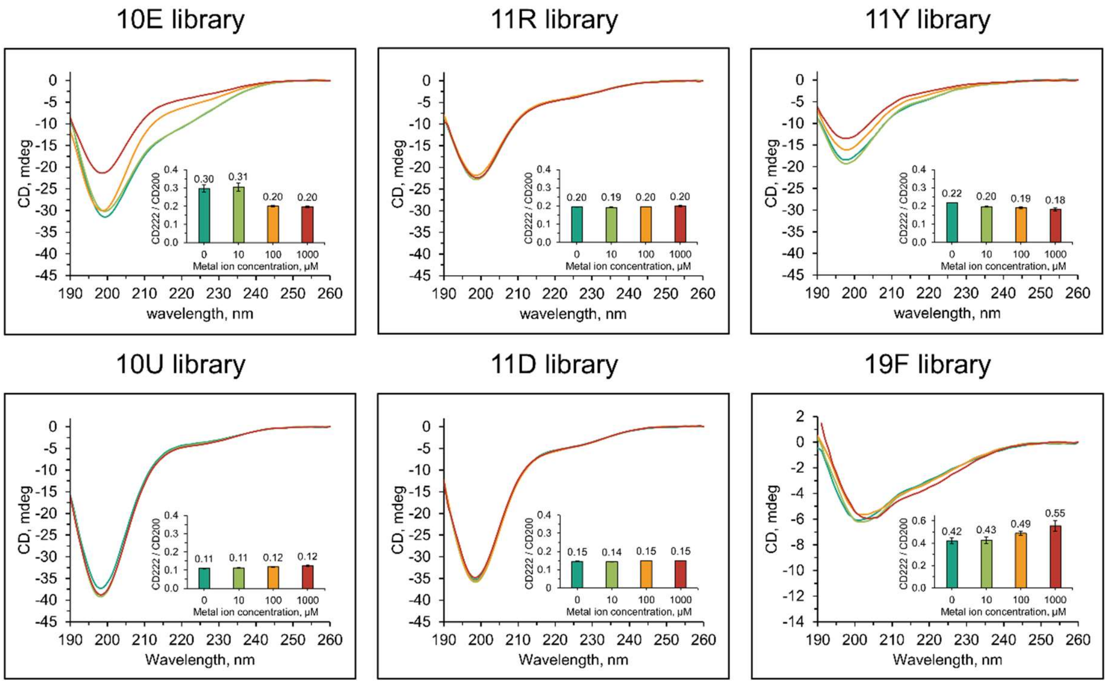

*Supplementary Figure S6. The effect of metal ions on the secondary structure of 25-mer combinatorial peptide libraries. CD spectra were measured for 0.2 mg/ml nominal peptide library solutions in a series of 10 mM Tris buffers at pH 7.4 supplemented with 0 (green line); 10 (yellow line); 100 (orange line); and 1,000 (red line) μM mixture of NaCl, KCl, $MgCl_2$, $MnCl_2$, and $ZnCl_2$. The inlet graph shows the ratios of CD signal at 222 to 200 nm.*
